# Non-Langmuir kinetics of DNA surface hybridization

**DOI:** 10.1101/2020.02.27.968081

**Authors:** L. Vanjur, T. Carzaniga, L. Casiraghi, M. Chiari, G. Zanchetta, M. Buscaglia

## Abstract

Hybridization of complementary single strands of DNA represents a very effective natural molecular recognition process widely exploited for diagnostic, biotechnology and nanotechnology applications. A common approach relies on the immobilization on a surface of single stranded DNA probes that bind complementary targets in solution. However, despite the deep knowledge on DNA interactions in bulk solution, the modelling of the same interactions on a surface are still challenging and perceived as strongly system-dependent. Here we show that a two dimensional analysis of the kinetics of hybridization, performed at different target concentration and probe surface density by a label-free optical biosensor, reveals peculiar features inconsistent with an ideal Langmuir-like behaviour. We propose a simple non-Langmuir kinetic model accounting for an enhanced electrostatic repulsion originating from the surface immobilization of nucleic acids and for steric hindrance close to full hybridization of the surface probes. The analysis of the kinetic data by the model enables to quantify the repulsive potential at the surface, as well as to retrieve the kinetic parameters of isolated probes. We show that the strength and the kinetics of hybridization at large probe density can be improved by a 3D immobilization strategy of probe strands with a double stranded linker.

**Statement of Significance:** Hybridization of nucleic acids strands with complementary sequences is a fundamental biological process and is also widely exploited for diagnostic purposes. Despite the availability of effective models for the equilibrium strength of freely diffusing strands, a general predictive model for surface hybridization is still missing. Moreover, the kinetics of hybridization is not fully understood neither in solution nor on a surface. In this work we show that the analysis of the kinetics of hybridization on a surface reveals and enables to quantify two main additional contributions: electrostatic repulsion and steric hindrance. These are general effects expected to occur not only on a surface but in any condition with large density of nucleic acids, comparable to that of the cellular nucleus.

## 1. Introduction

The formation of double-stranded DNA (dsDNA) from two complementary strands, called hybridization, is a fundamental process underlying DNA microarray technology (1), as well as the rapidly expanding field of DNA nanotechnology (2). DNA microarrays (DNA chips) have proven to be a powerful tool in many biomedical applications from detecting single-nucleotide polymorphisms to gene expression analysis (3). DNA chips are comprised of single stranded DNA (ssDNA) immobilized on a surface and acting as probes for complementary ssDNA in solution. Current research efforts in this field focus on two main goals: the development of high performance sensors and the design of molecular mechanisms to enhance the sensitivity and specificity of probe-target recognition (4)(5). In particular, DNA nanotechnology offers the opportunity to control the structure and function of complex supra-molecular systems and enables the design of programmable molecular machines (6).

Current limitation on the integration of DNA nano-machines on a biosensor surface is that the hybridization with a complementary strand immobilized on a surface generally displays a reduced affinity in comparison to the case in which both strands are freely diffusing in solution (7)(8). Interestingly, such difference between bulk and surface interactions is typically not observed for protein-protein interactions (e.g. antibody-antigen) and appears to be a characteristic of the nucleic acids (NA) recognition process. Different possible causes of this phenomenon have been proposed (9)(10)(11). More generally, the electrostatic repulsion plays an important role in the decreased hybridization strength on a surface. Indeed, ssDNA is a polyelectrolyte, in which each repeating unit bears one negative charge. The accumulation of ssDNA probes on a surface has been reported to induce an effective repulsive potential on freely diffusing complementary strands (12)(13), which shifts the equilibrium of hybridization in comparison to the same interaction in solution.

Similarly to equilibrium parameters, also the kinetics of surface hybridization significantly differs from the same process in the bulk (14)(15)(16)(17). Despite being fundamental to understand the origin of the equilibrium features, the kinetics of surface bound DNA hybridization is still poorly understood (18). A direct access to real-time binding curves without interference from labelling moieties is provided by label-free biosensors. Since the first studies performed by surface plasmon resonance (SPR) it has been shown that the realtime binding curves for DNA hybridization can depend on a number of factors, including the probe surface density, the probe distance from the surface, the presence of mismatches, and it can display non-exponential behaviour, in contrast with a simple Langmuir interaction model describing independent binding events (19) (14) (20). However, a general molecular interaction model to account for the kinetic curves for DNA hybridization on a surface is still missing. Indeed, the kinetics of hybridization is not fully understood even in the more standard case in which both complementary strands are freely diffusing in solution (21)(18). In this context, label-free biosensors not only represent a promising application field where to exploit DNA nanotechnology but they also provide an effective analytical tool to characterize DNA hybridization at molecular level. Several label-free biosensors have been exploited for sequence detection or quantification (22)(23)(24)(25)(26). Among these, Reflective Phantom Interface (RPI) has been demonstrated as a sensitive tool to characterize the kinetic and equilibrium parameters of biomolecular recognition process (27)(28) and, in particular, of fully or partially complementary oligonucleotides (11).

Here we show that the DNA hybridization kinetic curves acquired by label-free optical signal display marked deviations from a Langmuir behaviour in a wide range of conditions. We explored different surface density of complementary probes immobilized with or without a DNA linker, either ss or ds. We studied the hybridization at different concentration of target strand in solution and ionic strength. We found that both the equilibrium behaviour and the kinetics of hybridization show discrepancies from an ideal Langmuir interaction in all explored conditions. The results support the primary effect of electrostatic repulsion originating in proximity of the surface due to NA accumulation. Moreover, close to saturation of the surface probes by complementary targets we observed a marked decrease of the apparent kinetic constant for hybridization as a consequence of surface crowding. The measured reduction of hybridization affinity at large local NA concentrations strongly affects the results of DNA or RNA microarrays and biosensors and can play a biological role in the cellular environments rich of DNA, such as the nucleus. In general, the enhanced repulsion observed for the hybridization at large DNA local density could contribute to keep a large specificity of pairing even in a DNA crowded environment.

## 2. Materials and methods

### 2.1 DNA strands and reagents

We studied the kinetics of hybridization of a 12mer model sequence with different surface-immobilized complementary probes. As schematically shown in Figure 1, the simplest interaction with a 12mer probe (no linker) was compared to that measured with probes having additional ss-linker or ds-linker. The NA sequences used in this work are reported in Table 1. Probe strands p1 and p2 were immobilized on the RPI sensing surface and t1 was used as complementary target strands in solution. Strand cp2 was optionally used to make a dsDNA spacer at the base of p2. ssDNA were purchased from Integrated DNA Technologies (Leuven, Belgium) with high-quality Ultramer synthesis. Strands p1 and p2 were amine-modified at C6 carbon of 5’ terminal (5AmMC6 in Table 1). The surface immobilization of amine-terminated ssDNA was achieved by coating the RPI sensing chip with MCP2 or MCP4 copolymers from Lucidant Polymers (Sunnyvale, CA, USA). They are copolymers of dimethylacrylamide (DMA), N-Acryloyloxysuccinimide (NAS), and 3- (Trimethoxysilyl)propyl methacrylate (MAPS) and they differ only in the co-monomer molar ratio: 97:2:1 in MCP2 and 89:10:1 in MCP-4. The fraction of amine-reactive sites of MCP4 is five times larger than that of MCP2. All buffers and reagents were purchased from Sigma-Aldrich and prepared according to common protocols using Milli-Q pure water.

**Table 1.**
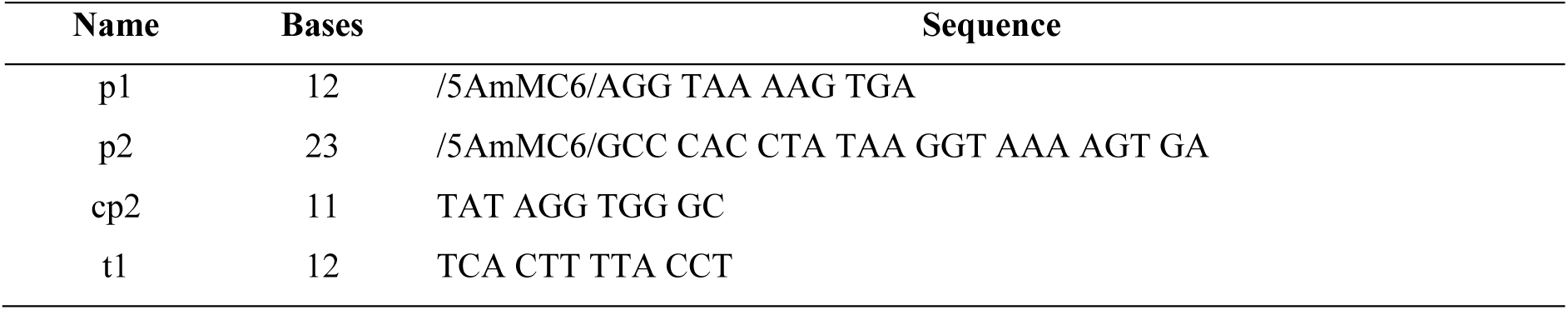
DNA sequences.

**Figure 1.**
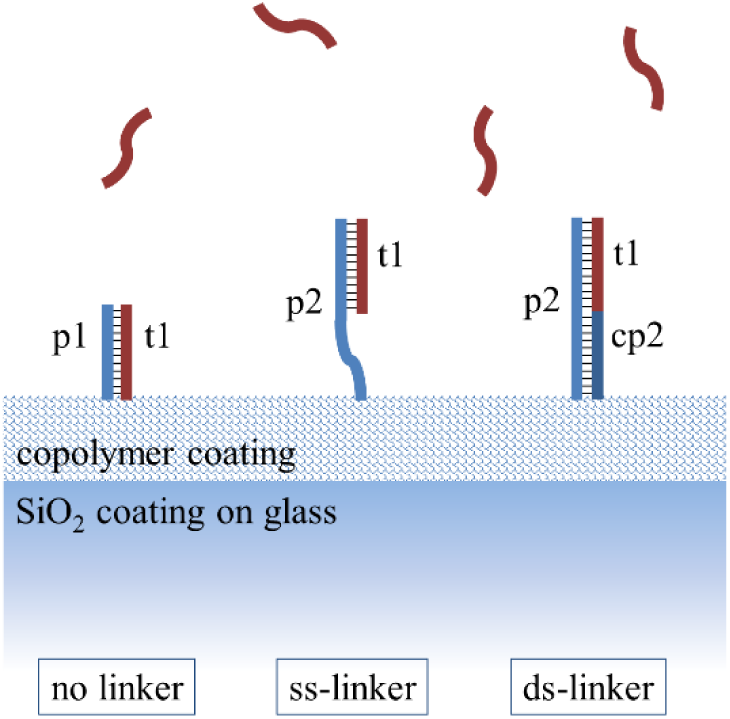
Schematics of surface probe types. The 12-mer target strand t1 (dark red) binds to a complementary strand probe (blue) immobilized on the RPI sensing surface by a 3D copolymer coating. Three types of DNA probes were investigated: a complementary 12-mer strand (p1, no linker scheme) and longer probes formed by a ss strand (p2, ss-linker) or a ds strand (p2 + cp2, ds-linker) terminated with the complementary sequence.

### 2.2 RPI sensor preparation and measurement

DNA probe strands were covalently immobilized on the surface of RPI sensing chips in spots with 150-200 μm diameter following the procedure described in (28). Briefly, 8 × 12 mm wedge-like chips of F2 optical glass (Schott, Mainz, Germany) coated with an anti-reflection layer of SiO_2_ were plasma cleaned and dip-coated with MCP2 or MCP4 copolymer (29). Droplets of spotting buffer (Na_2_HPO_4_ pH 8.5 150 mM) containing amine-terminated DNA probes at concentrations from 1 μM up to 30 μM were deposited on the chips surface by an automated noncontact dispensing system (sciFLEXARRAYER S5, Scienion AG, Berlin, Germany). After overnight incubation, the chip surface was rinsed with blocking buffer (Tris HCl pH 8 10 mM, NaCl 150 mM, ethanolamine 50 mM) and distilled water and then dried. The sensor cartridges were prepared by gluing the glass chips on the inner wall of 1-cm plastic cuvettes. The cartridges were stored at 4°C before use. Target ssDNA strand t1 and strand cp2 were suspended before use in measuring buffer (10 mM Tris-HCl, 0.02% NaN_3_, pH 8.0 + NaCl at different concentrations depending on the measurement).

The RPI measurements were performed by using the apparatus and the analysis algorithm described in (28). The sensor cartridges were filled with 1.3 ml of measuring buffer. The ionic strength was adjusted by adding NaCl from 75 mM up to 220 mM. The cartridges were kept at 23 °C during the measurement through a thermalized holder and rapid mixing of the solution was provided by a magnetic stirring bar. Sample spikes of target ssDNA were performed by adding 50 μl of measuring buffer containing different amounts of target molecules to a final concentration in the cartridge from 0.5 nM up to about 1.5 μM. Time sequences of RPI images of the spotted surface were analysed by a custom Matlab program (Mathworks, Natick, MA, USA) in order to obtain the brightness of each spot as a function of time *t* and convert it into the total mass surface density of molecules *σ*(*t*) and in the mass surface density of the target molecules *σ*_*t*_(*t*) = *σ*(*t*) – *σ*_*p*_(*t*), where *σ*_*p*_(*t*) is the mass surface density of immobilized probe molecules measured before the addition of the target ssDNA in solution. The analysis of the hybridization curves was performed on *σ*_*t*_(*t*) traces obtained by averaging at least 6 spots with identical composition. The number surface density of probe, *s*_*p*_, and target molecules, *s*_*t*_, was obtained dividing *σ*_*p*_ and *σ*_*t*_ by the corresponding molecular mass, respectively.

### 2.3 Analysis of surface hybridization by Langmuir model

The hybridization kinetics curves *σ*_*t*_(*t*) measured by RPI were analysed either assuming a standard Langmuir model (15)(38) or the non-Langmuir kinetic model described in section 3.2. The main assumptions of the Langmuir model are that the surface provides a finite number of independent binding sites (probes) holding at most one target molecule each, the binding sites are all equivalent, their properties do not change during the binding process, and there are no interactions between target molecules bound on adjacent sites. Under these assumptions, the time evolution of the fraction of hybridized surface probes *ϕ*(*t*) is given by

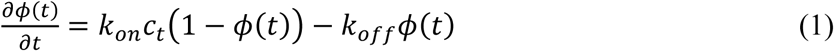

where *c*_*t*_ is the concentration of target ssDNA in solution and *k*_*on*_ and *k*_*off*_ are the kinetic rate constants for association (hybridization) and dissociation, respectively. For a concentration jump to *c*_*t*_ at *t* = 0, the solutions of Eq. 1 are exponential growth functions with the form

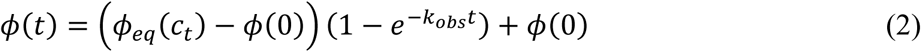

where

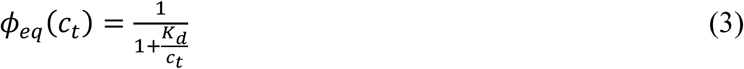

is the equilibrium plateau value, which depends on the dissociation equilibrium constant *K*_*d*_ = *k*_*off*_ /*k*_*on*_ of probe-target hybridization, and

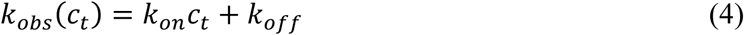

is the observed hybridization rate. The mass surface density, *σ*_*t*_(*t*), or the number surface density of target, *s*_*t*_(*t*), at a given time *t* after an increase of concentration *c*_*t*_ and the asymptotic equilibrium values *σ*_*eq*_ or *s*_*eq*_ are given by *σ*_*t*_(*t*) = *ϕ*(*t*)*σ*_*∞*_ and *σ*_*eq*_ = *ϕ*_*eq*_*σ*_*∞*_ or by *s*_*t*_(*t*) = *ϕ*(*t*)*s*_*∞*_ and *s*_*eq*_ = *ϕ*_*eq*_*s*_*∞*_, respectively, where *σ*_*∞*_ and *s*_*∞*_ are the mass surface density and the number surfaces density of target at saturation reached at large *c*_*t*_.

## 3 Results

### 3.1 Effect of probe surface density on strength and kinetics of hybridization

We studied the kinetics of the hybridization process of ssDNA oligomers in solution (targets) with complementary strands (probes) immobilized on the surface of the RPI label-free biosensor. We focused on target oligomers with a length of 12 bases because they are long enough to provide rather large hybridization strengths and large label-free signals, and small enough to observe a clear dependence of their interaction parameters on different experimental conditions. We explored both the equilibrium constant and the kinetic rate constant for complementary probes with no additional linker or with a ss or ds linker strand, as shown in Figure 1. The injection into the RPI measuring cell of target ssDNA provides an increase of signal corresponding to the surface density of targets *σ*_*t*_(*t*) binding to the immobilized probes. Figure 2 reports label-free hybridization curves measured for probes with no linker (probe p1 in Figure 1) after the addition of targets in solution at the concentration *c*_*t*_ of 100 nM. The curves correspond to different spot families on the same RPI chip produced with different probe concentrations in the spotting buffer, from 2.5 µM up to 30 µM. All curves reached a stable asymptotic value of target surface density *σ*_*eq*_ at long time. However, both the asymptotic amplitude and the time required to reach such asymptotic value depend on the spotting concentration of probes. As other label-free biosensors, the RPI DNA sensor enables a direct measure of the mass surface density of probes *σ*_*p*_ from the brightness of the spots before the addition of target in solution. The number of captured target strands is roughly proportional to the number of surface probes, although it remains smaller (Figure S1, Supporting Information), indicating that a fraction of probe strands on the surface are not accessible to the target. In the experiment reported in Figure 2, the hybridization yield *Ψ*, that is the fraction of surface probes hybridized with the target, was about 30%. More generally, considering all the measurements reported in this work, the obtained *Ψ* was overall 50% ± 20%, with a tendency of copolymer coating MCP4 to provide values of *Ψ* slightly larger than MCP2.

**Figure 2.**
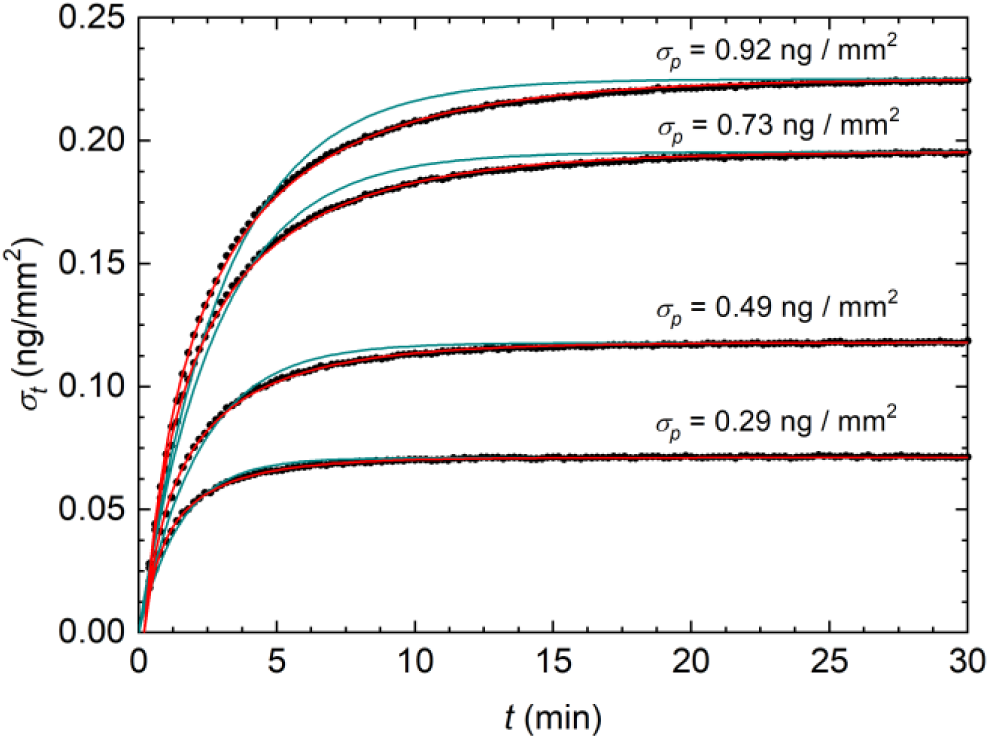
Hybridization kinetic curves measured by RPI. (a) Binding curves (black dots) expressed as mass surface density measured after the injection of 100 nM of target DNA in solution with 150 mM NaCl. The different curves refer to spots on the same RPI sensor with different surface density *s*_*p*_ of DNA probes (no linker type), as reported in the Figure. The DNA probes are immobilized via MCP2 copolymer. The continuous lines represent the fits with single exponential growth functions (blue) and numerical solutions of Eq. 6 (red).

The binding curves reported in Figure 2 also show a marked dependence of the hybridization kinetics on the surface density of probes. Smaller probe densities not only yield smaller amplitudes but also shorter times to reach the equilibrium. Under the hypothesis of an ideal interaction described by the Langmuir model, we fitted the hybridization curves with simple exponential growth functions (Eq. 2), from which we obtained the characteristic rate *k*_*obs*_. As shown in Figure 2, only the binding curve corresponding to the spots with the smallest *σ*_*p*_ was well fitted by an exponential growth (blue curves), and the deviation progressively increases with increasing *σ*_*p*_. This behaviour suggests that the Langmuir interaction model does not represent well the hybridization kinetics between 12mers for large surface densities of probes. Binding curves that are not well fitted by single exponential growth curves are commonly observed by label-free biosensors and their interpretation typically involves different causes, including heterogeneity of the surface binding sites or conformational changes of probes and targets (30). A general approach is based on the assumptions of multiple Langmuir-like processes with different kinetics that sum up and yield multi-exponential binding curves (31). Here we adopted a different strategy based on a deeper investigation of the scaling of the amplitudes and rates of the binding curves progressively increasing the concentrations *c*_*t*_ of target in solution.

We performed sequential additions of target strands in solution, obtaining a target concentration *c*_*t*_ from 0.5 nM up to 1562.5 nM in the same RPI cell, and we measured realtime hybridization curves for spot families with different spotting concentration of probes, hence obtaining a matrix of binding curves for different *c*_*t*_ and *σ*_*p*_, as shown in Figure 3a. The inspection of the data at intermediate target concentrations (central column of plots in Figure 3a) shows that the effect of *σ*_*p*_ on the amplitude and rates of the hybridization curves is qualitatively similar to that reported in Figure 2. Remarkably, in this case all the measured hybridization curves are well fitted by exponential growth functions (black curves), even at large *σ*_*p*_, because the dynamic range of each individual curve *σ*_*t*_(*t*) is typically smaller. This is equivalent to observing only a portion of the full curve *σ*_*t*_(*t*) from *σ*_*t*_(*0*) = 0 to *σ*_*eq*_(*c*_*t*_), as those in Figure 2. The fit of the measured *σ*_*t*_(*t*) curves with exponential functions enables to extract amplitudes and rates as a function of *c*_*t*_ and for various values of *σ*_*p*_. In this way, the matrix of binding curves of Figure 3a was converted into two matrices, one for the asymptotic amplitudes *σ*_*eq*_(*c*_*t*_,*σ*_*p*_) and the other for the hybridization rates *k*_*obs*_(*c*_*t*_,*σ*_*p*_). The results are reported as plots at constant *σ*_*p*_ in Figure 3b and 3c (blue squares). All measured *σ*_*eq*_(*c*_*t*_) (Figure 3b) can be approximately fitted with a simple Langmuir model, according to Eq. 3 (continuous blue curves). The corresponding equilibrium dissociation constant *K*_*d*_, indicated by the dashed lines in the figures, increases with the spotting concentration of probes, suggesting a weakening of hybridization strength with the increase of *σ*_*p*_. However, at a closer inspection of the amplitude data a small systematic deviation from the ideal behaviour can be observed: the fit tends to slightly underestimate the data points at concentrations *c*_*t*_ smaller than *K*_*d*_, and overestimate those for *c*_*t*_ larger than *K*_*d*_. For what concerns the measured hybridization rates *k*_*obs*_(*c*_*t*_) (Figure 3c, blue squares and continuous curve), the analysis shows that the expected linear dependence on *c*_*t*_ (Eq. 4) is confirmed up to about *c*_*t*_ = 100 nM. The intercept of *k*_*obs*_(*c*_*t*_ → 0), corresponding to *k*_*off*_, appears to be constant and independent on *σ*_*p*_, whereas the slope, corresponding to *k*_*on*_, decreases with *σ*_*p*_.

**Figure 3.**
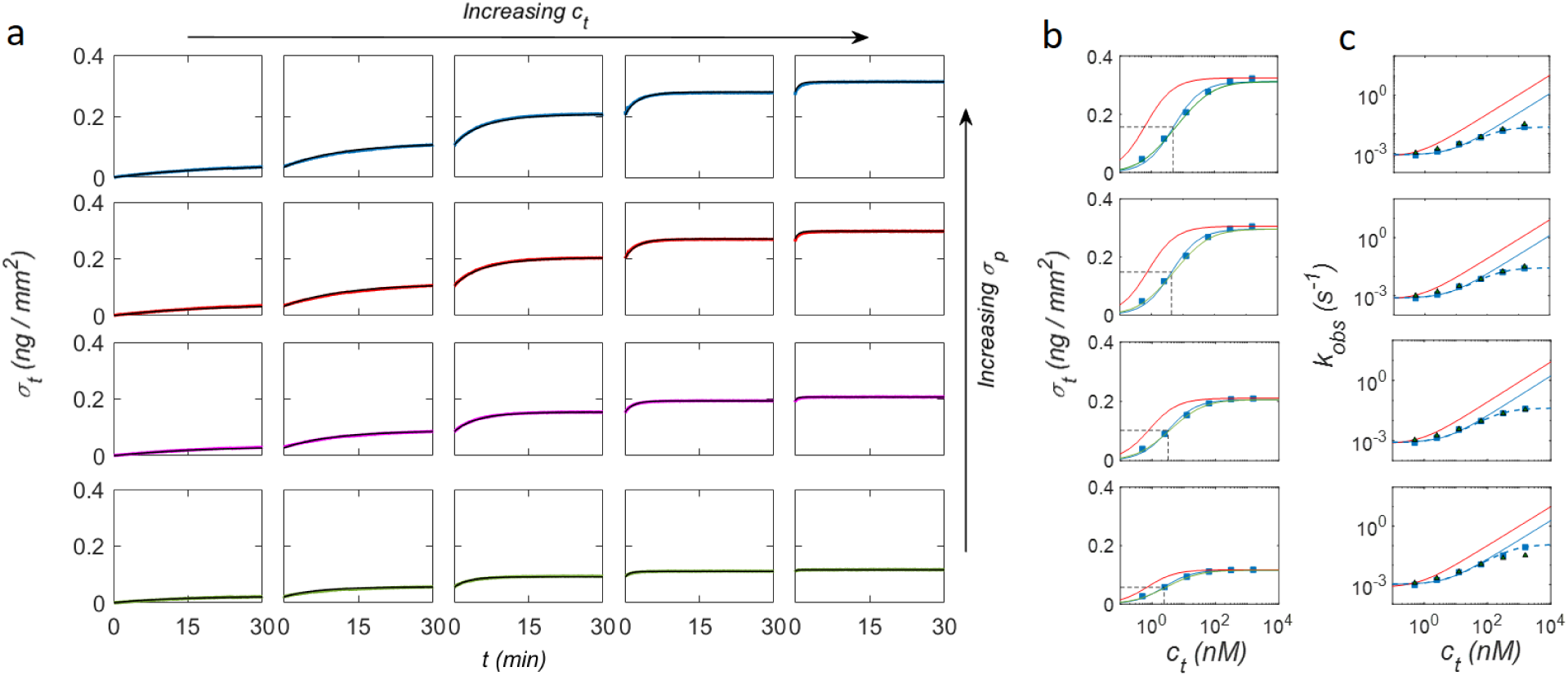
Hybridization kinetic curves at different target concentration and probe surface density. (a) RPI binding curves measured on the same sensor on spots with probe density 0.19 ng/mm^2^ (green), 0.24 ng/mm^2^ (purple), 0.38 ng/mm^2^ (red) and 0.41 ng/mm^2^ (cyan), for increasing concentrations of target strand in solution: (from left to right) 0.5 nM, 2.5 nM, 12.5 nM, 62.5 nM and 312.5 nM. The black curves are fits to the data by single exponential growth functions. (b) Equilibrium asymptotic amplitudes and (c) kinetic rates obtained from exponential fits of the hybridization curves of panel a (blue squares). The blue lines are fits with amplitudes and rates obtained from a Langmuir model. The last two points at the largest concentrations are excluded from the fit of the rates. The green lines and the green triangles are the values obtained from the fit with the NLER model. The red lines represent the Langmuir behaviour extrapolated from the NLER fit for *Γ* = 0.

The observed rates clearly deviate from the ideal linear dependence on *c*_*t*_ only for the largest concentrations of target, when the fraction of hybridized active probes *ϕ* is close to 1. Figure 3c shows that the rates measured at *c*_*t*_ > 100 nM are smaller than the values extrapolated from the dependence of *k*_*obs*_(*c*_*t*_) at smaller *c*_*t*_, and the deviation from the linear scaling with *c*_*t*_ is progressively more pronounced at increasing *σ*_*p*_. We assumed that, close to saturation, the remaining small fraction of available single stranded probes yield to a slower association kinetics, possibly because of their close proximity to other single stranded probes or hybridized duplexes (32). This interpretation is consistent with the larger deviations from ideal linear scaling observed for larger *σ*_*p*_, hence for smaller average probe-probe distance, and is also consistent with the absence of this effect for the ds-linker probe type, in which a larger distance among neighbourhood probes is maintained by the larger volume and stiffness of the ds segment. It is worth noting that the inhomogeneous probe-probe distance obtained by random immobilization of DNA strands has been also proposed as the cause of the shape of the melting curves for surface-immobilized DNA (33).

To empirically describe the observed reduction of the apparent *k*_*on*_ at large *c*_*t*_, we assumed a characteristic value of the fraction of hybridized probes *ϕ* = *ϕ\**, at which this phenomenon occurs. In order to estimate *ϕ\**, the observed rate *k*_*obs*_(*c*_*t*_) was fitted in the full range of concentrations with the following equation:

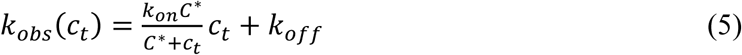

where the parameter *C\** represents the target concentration at which the apparent *k*_*on*_ displays a twofold decrease relative to the low concentration value. Eq. 5 well fits the data reported in Figure 3c (dashed lines). We converted *C\** into the corresponding values of *ϕ\** through Eq. 3. Figure 4a shows that *ϕ\** decreases as a function of the saturation value of the number surface density of target strands *s*_*∞*_ for the no linker and ss-linker hybridization types. The observed behaviour is consistent with the interpretation of *ϕ\** as the fraction of probe strands with a large enough distance from each other on the surface to grant free accessibility to the target strand. In each conditions, an average fraction *ϕ\** of probes display a kinetics of hybridization unaffected by surface crowding. Therefore, this effect is not expected to affect the hybridization parameters for concentrations much lower than *C\**.

**Figure 4.**
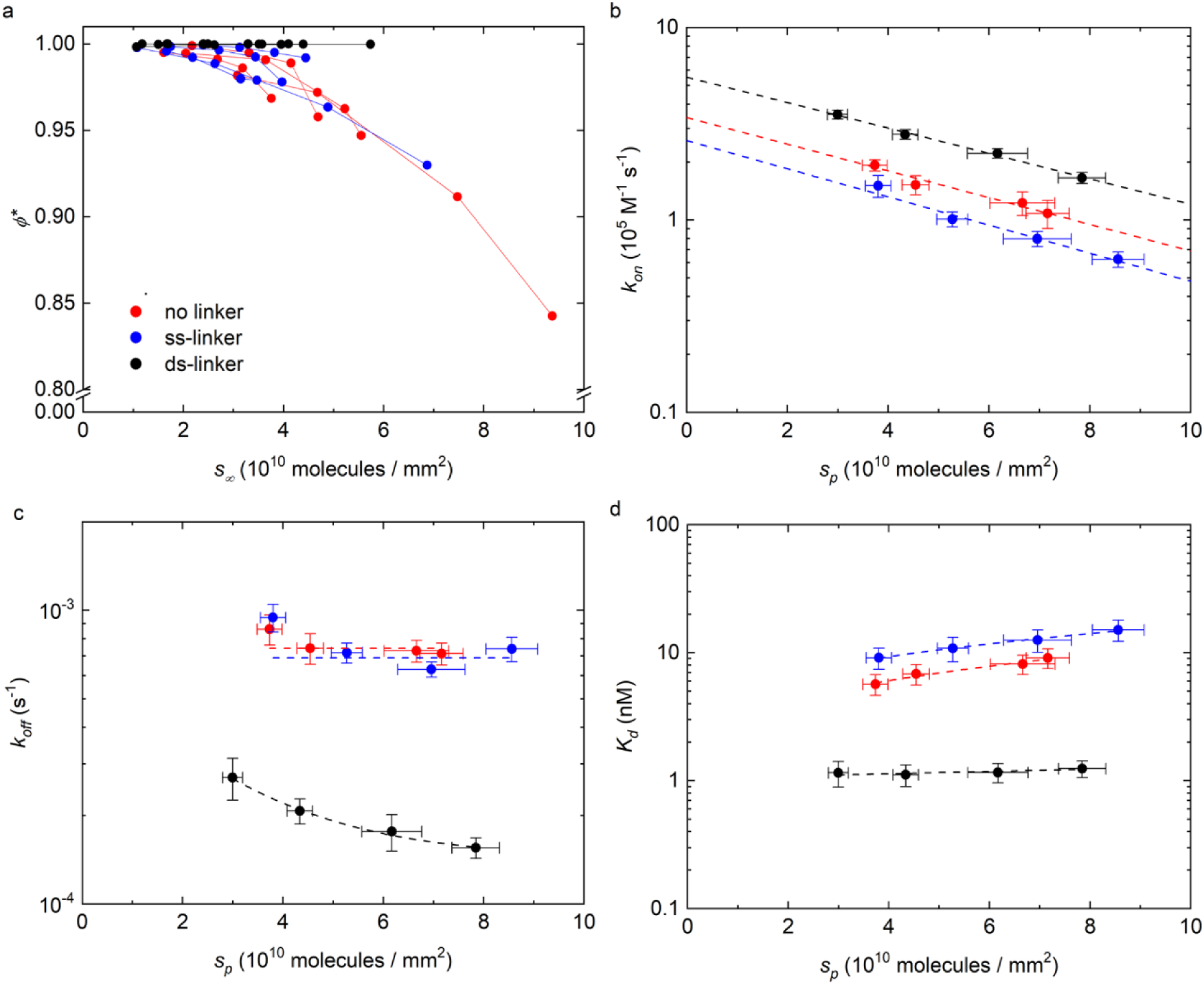
Dependence on DNA surface density of the equilibrium and kinetic parameters for hybridization. (a) Fraction of hybridized active probes *ϕ\** as a function of target surface density *s*_*∞*_ at saturation for all four experiments and all probe types. Kinetic rate for association (b) and dissociation (c) obtained for different probe types. The dashed lines represent fits to the data with the same colour: constant values (panel c, red and blue) or exponential decays (all the curves of panel b and black curve in panel c). (d) Dissociation equilibrium constant for different probe types. The dashed lines are linear fits shown to guide the eye. In panel b, c and d the data points are average values of four experiments and the error bars are the standard deviations. In all panels, the colours refer to different probe types, as indicated in panel a.

The analysis of the amplitude and rate data of Figure 3b and 3c by a simple Langmuir model (Eq. 3 and 5) enables to quantify the interaction parameters for the hybridization at different values of the number density of probes *s*_*p*_. We obtained the kinetic rate constant for association, *k*_*on*_, and dissociation, *k*_*off*_, by a global fit of the amplitude and rate dependence on *c*_*t*_ with the constraint *K*_*d*_ = *k*_*off*_/*k*_*on*_. We repeated the analysis for the different probe types sketched in Figure 1 and for the two copolymer coatings, MCP2 and MCP4. No appreciable difference was observed for the hybridization kinetics measured from the two coatings at similar *s*_*p*_. However, the use of both coating types enabled to slightly extend the overall range of *s*_*p*_. As reported in Figure 4b and 4c, we found similar values of the kinetic constants for no linker and ss-linker probes and larger *k*_*on*_ and much smaller *k*_*off*_ for ds-linker. We also observed a systematic decrease of *k*_*on*_ on increasing the surface density *s*_*p*_ for all probe types. In contrast, *k*_*off*_ is nearly constant in the case of no linker and ss-linker probes. Therefore, the increase of *K*_*d*_ with *s*_*p*_, reported in Figure 4d, primarily results from *k*_*on*_, in these cases. Differently, for the ds-linker probes we obtained much smaller values of *K*_*d*_, hence a stronger hybridization strength, weakly dependent on *s*_*p*_. We ascribed the peculiar behaviour of the ds-linker probes primarily to the presence of the additional base stacking interaction due to the double strand adjoining the probe sequence, which can be as large as 1.5 kcal/mol in the considered experimental conditions (34) (35). In contrast, the observed decrease of *k*_*on*_ with *s*_*p*_ reported in Figure 4b is primarily ascribed to an additional effect originating from the electrostatic repulsion between NA strands, as discussed below.

### 3.2 Non-Langmuir kinetic model with electrostatic repulsion

The interaction between a target ssDNA and its complementary strand immobilized on a surface is known to be affected by electrostatic repulsion (12)(13)(17). In particular, this effect is expected to increase with the overall surface density of NA. Consequently, the mean electrostatic repulsion in the proximity of the surface can increase during the hybridization, which brings more NAs, hence more charges, onto the surface. This condition yields to an apparent reduction of the hybridization strength at equilibrium, which depends on the fraction of hybridized probes on the surface. Therefore, the hybridization process could show deviations from a simple Langmuir model even at small target concentrations *c*_*t*_ and fractional coverage of active probes *ϕ*. A simple theoretical solution of the equilibrium condition has been proposed by Vainrub and Pettitt (VP) (36) introducing a mean-field free-energy penalty for hybridization proportional to the surface fraction of bound active probes *ϕ* and accounting for an effective electrostatic repulsive potential confined in a thin surface layer. The model has been further refined by Halperin, Buhot and Zhulina (HBZ) (37) allowing for a variable thickness of the repulsive layer, hence describing the hybridization also at low ionic strength. The notion of a repulsive potential originating at the surface of DNA biosensors and DNA arrays enables to compute more accurate equilibrium solutions for the hybridization process (9) (12)(38). In contrast, an effective general model to account for the measured kinetics of hybridization is still missing. An influence of the surface probe density on the kinetics of DNA hybridization has been often observed in biosensor measurements (14) and a few studies have proposed theoretical frameworks accounting for electrostatic repulsion (32)(17).

On the basis of the VP and HBZ equilibrium models and of the previous studies on kinetics modelling we developed a simple approach to account for the effect of a repulsive potential in the proximity of the probe layer on the kinetics of hybridization. Figure 5a shows a schematic representation of the model: the accumulation of negative net charge on the surface yields to a repulsive electrostatic potential, which, in a simple approximation, we assume to have a step-like profile with a characteristic thickness *h*. At a distance larger than *h* from the surface, the potential is that of the bulk solution. The model also comprises the notion of a dissociation constant *k*_*off*_ substantially independent from the probe surface density, as suggested by the experiments shown in Figures 3 and 4. Under these assumptions, the time evolution of the surface fraction of hybridized probes, for *ϕ* < *ϕ\**, is described by

**Figure 5.**
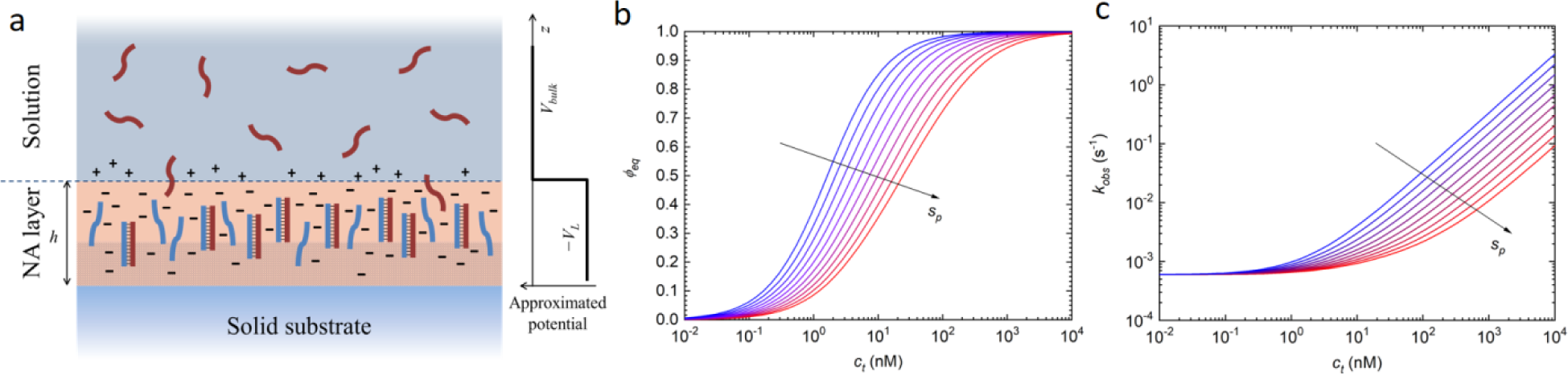
Schematic representation and numerical solution of the NLER model. (a) The surface region is rich in NA probes (blue) immobilized on the copolymer coating (shaded area) and hence provides a negative net charge, which further increases upon hybridization with the target strands (dark red). The electrostatic potential is approximated by a step function having a value lower than that of the bulk solution up to a distance *h* from the RPI solid surface. (a) Fraction of hybridized probes at equilibrium and (b) kinetic rate computed for different surface density of probes, from 10^10^ mm^-2^ (blue) to 10^11^ mm^-2^ (red). The curves were computed with fixed kinetic parameters 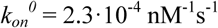 and *k*_*off*_ = 7.2·10^−4^ s^-1^ and for *Γ* = *γs*_*p*_, with *γ* = 2·10^−11^ mm^2^.

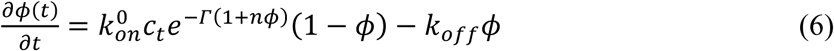

where 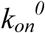 is the association kinetic rate in the ideal condition of negligible repulsive interaction, *n* is the ratio between the length of target and probe strands expressed in number of bases, and *Γ* represents the electrostatic penalty associated with entry of charged ssDNA target into the probe surface layer, as predicted in the VP and HBZ models. We named this kinetic model Non-Langmuir model with Electrostatic Repulsion (NLER). Eq. 6 differs from a Langmuir kinetic model (Eq. 1) only for the exponential term *e*^−*Γ*(1+*nϕ*)^, which accounts for the electrostatic repulsion experienced by the target strands in the proximity of the surface with the immobilized probes. Remarkably, this exponential term can be considered either as a correction coefficient applied to 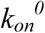, hence reducing the effective association time because of the repulsive free-energy barrier, or equivalently as a term applied to the concentration *c*_*t*_, hence reducing the amount of target DNA effectively entering the probe surface layer. Eq. 6 also indicates that the effect of the repulsive interaction yields a behaviour different from a simple Langmuir process for values of the product *Γ·n* close to or larger than 1. In this case, the surface density of charges changes significantly between the conditions of *ϕ* = 0 (only probe strands) and *ϕ* = 1 (all active probes hybridized with the target strands). Considering the probe schemes shown in Figure 1, the value of parameter *n* is 1 for the no linker type, 1/2 for the ss-linker and 1/3 for the ds-linker. Notably, even if the kinetics becomes indistinguishable from a Langmuir process for small *n*, the repulsive interaction can still be relevant if *Γ* is non-negligible, and both the observed association rate constant and the equilibrium constant effectively incorporate the term *e*^−*Γ*^.

Numerical solutions of Eq. 6 well describe the measured hybridization curves with a minimum set of parameters. In particular, Eq. 6 describes both the non-exponential shape of the hybridization kinetic curves for large *c*_*t*_ jumps, as shown in Figure 2 (red curves), and the dependencies of *ϕ*_*eq*_ and *k*_*obs*_ on *c*_*t*_, as those reported in Figure 3. Figure 5 shows that *ϕ*_*eq*_(*c*_*t*_) and *k*_*obs*_(*c*_*t*_) calculated from Eq. 6 differ from those obtained with the Langmuir model. The amplitudes *ϕ*_*eq*_(*c*_*t*_) of the simulated hybridization curves are shifted at larger *c*_*t*_ and increase with a smaller slope for larger values of *Γ*. Interestingly, a similar behaviour can be also obtained by standard general models accounting for a distribution of interactions with different *K*_*d*_ or by the widely used Sips isotherm (19)(39). Analogously, the observed rates *k*_*obs*_(*c*_*t*_) display a progressively weaker dependence on *c*_*t*_ for larger values of *Γ*.

We used the numerical solutions of Eq. 6 to perform a two-dimensional fit of the measured hybridization curves *σ*_*t*_(*t*) at different *c*_*t*_ and *s*_*p*_ (Figure 3a) in order to extract the value of *Γ*, 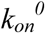 and *k*_*off*_. The green curves in Figure 3b and the green triangles in Figure 3c report the fit to *σ*_*eq*_(*c*_*t*_) and *k*_*obs*_(*c*_*t*_), respectively. As a comparison, the red curves in Figure 3b and 3c report the amplitudes and rates extrapolated to the absence of repulsive potential at the surface, hence for *Γ* = 0. In this ideal condition, Eq. 6 describes a Langmuir process with kinetic rates 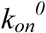 and *k*_*off*_ and thus with 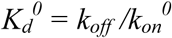. The behaviour of both the equilibrium amplitudes and the observed rates become increasingly non-Langmuir as the surface density of probes increases. The shift at larger concentrations either for the amplitude plots (Figure 3b) or the rates (Figure 3c) provides the value of the term *Γ* in these process. The values of *Γ* are consistent with a liner scaling with *s*_*p*_ (Figure S2, Supporting Information)(37). Therefore, we assumed

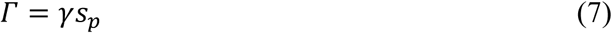

and, for each experiment, we fitted the amplitude and rate data at different *c*_*t*_ and *s*_*p*_ with only one value of 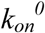 and one value of *γ*. In the case of the no linker and ss-linker probe type, also *k*_*off*_ was assumed to be independent on *s*_*p*_, whereas it was assumed to provide a linear dependence on *s*_*p*_ for the ds-linker probes. The average values of the kinetic parameters obtained from four experiments for each of the three probe types considered are reported in Table 2. The obtained kinetic rate constants are very similar for the no linker and the ss-linker probes and show a larger 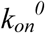 and a smaller *k*_*off*_ for the ds-linker case. The value of *Γ* at a standard surface density of *s*_*p*_, = 10^11^ mm^-2^ is of the order of 1 for all probe types and shows a minimum value for the ds-linker type. Indeed, the value of *Γ* is expected to primarily depend on different physico-chemical variables affecting the charge interactions between NAs. A deeper insight on this dependence is provided by the study of the hybridization at different ionic strengths.

**Table 2.** Measured parameters for DNA hybridization at 150 mM NaCl.

| Probe type | Probe sequence | $\Gamma^a$ | $K_d^0$ (nM) | $k_{on}^0$<br>($10^5 \text{ M}^{-1}\text{s}^{-1}$) | $k_{off}^0$<br>($10^{-4} \text{ s}^{-1}$) |
| --- | --- | --- | --- | --- | --- |
| no linker | p1 | $1.2 \pm 0.1$ | $2.2 \pm 0.6$ | $3.7 \pm 0.7$ | $7.7 \pm 0.6$ |
| ss-linker | p2 | $1.3 \pm 0.3$ | $3.0 \pm 1.0$ | $3.0 \pm 1.0$ | $6.5 \pm 0.8$ |
| ds-linker | p2 + cp2 | $0.8 \pm 0.3$ | $0.3 \pm 0.1$ | $6.4 \pm 0.7$ | $1.7 \pm 0.5$ |

<sup>a</sup> obtained for $s_p = 10^{11} \text{ mm}^{-2}$

### 3.3 Effect of ionic strength on the hybridization kinetics

The role of ionic interactions can be in general modulated by changing the solution concentration of salt, which provides the counter ions that screen the chain ions. In particular, the hybridization can be partially or totally inhibited at large surface densities of probes and low concentrations of salt in solution (13). To quantitatively account for the influence of ionic strength on binding kinetics we investigated the hybridization curves for solutions containing different salt concentration, ranging from below to above the value of ionic strength *I*_*s*_ = 150 mM, which best approximates the physiological conditions. In general, we observed an increase of the hybridization rates with *I*_*s*_ and such dependence is more pronounced at low surface density of probes. Figure 6 reports the observed hybridization rates *k*_*obs*_ as a function of *s*_*p*_ obtained by exponential fits of the hybridization curves *σ*_*t*_(*t*) measured for different ionic strengths at the same target concentration *c*_*t*_ = 12.5 nM. For all salt concentrations, the measured rates constantly decrease with *s*_*p*_ and tend to converge to similar values at large *s*_*p*_. In the explored regimes, the values of *k*_*obs*_ span about one order of magnitude, from the smallest values measured at large probe density and small salt concentrations up to those extrapolated for small *s*_*p*_ and large *I*_*s*_. This confirms that the hybridization kinetics can be controlled by either the surface density of probes or the salt concentration.

**Figure 6.**
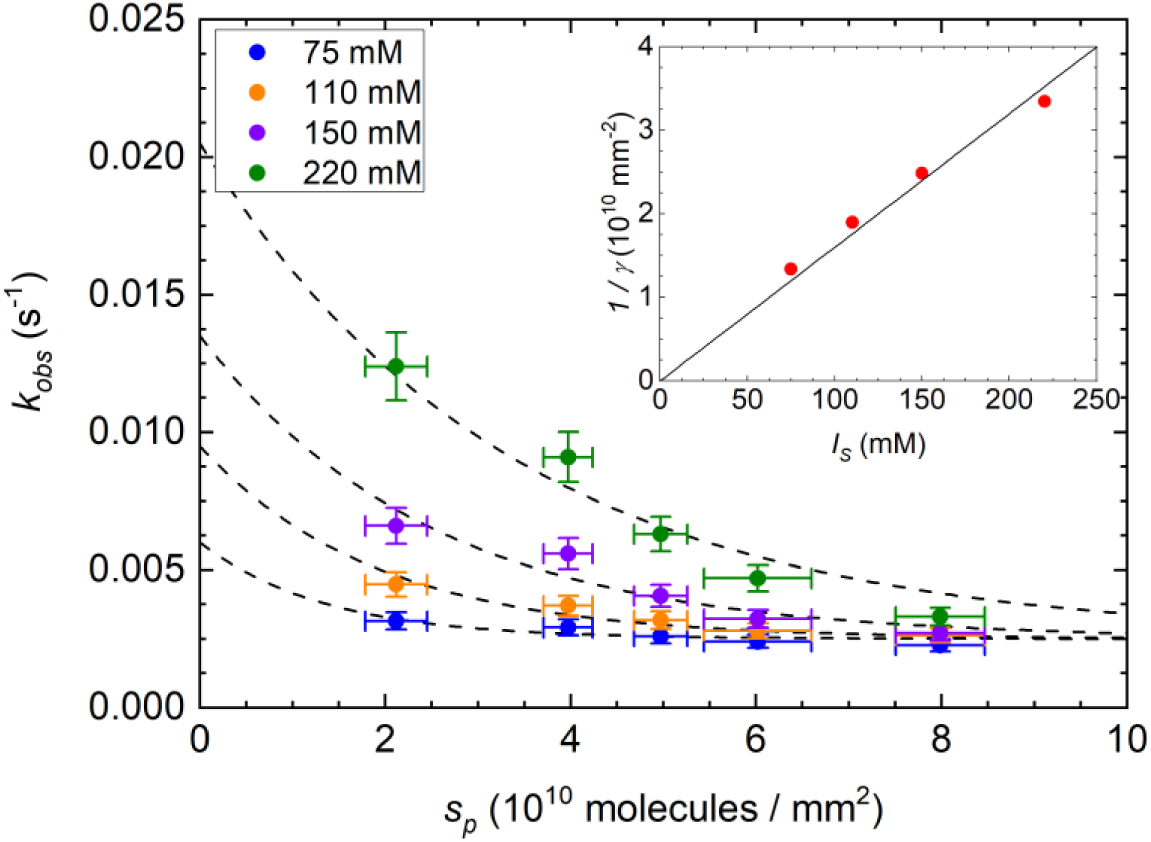
Dependence of the measured kinetic rates for hybridization on the ionic strength. RPI binding curves measured at target concentration of 10 nM, for different surface density of probes (no linker type) and for different ionic strength, as indicated in the figure legend. The vertical error bars represent the standard deviation of observed rates calculated from three experiments at 150mM NaCl. The dashed curves are fits with exponential decay functions, constrained to the same asymptotic value at large *s*_*p*_ and to an initial value at *s*_*p*_ = 0 linearly increasing with the ionic strength. Inset: scaling of the characteristic surface *γ* with the ionic strength obtained from the exponential decay fit of *k*_*obs*_ (red dots), and linear fit with slope 16·10^−10^ mm^-2^/M (black line).

The observed behaviour of *k*_*obs*_ as a function of surface probe density at different *I*_*s*_ is compatible with Eq. 6. The data reported in Figure 6 were fitted with curves *k*_*obs*_(*s*_*p*_) obtained for Eq. 6 assuming a linear dependence of 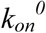 as a function of *I*_*s*_. Considering the range of salt concentrations explored in this study, the linear dependence of 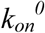 is consistent with previous measurements of kinetic rate constants for hybridization of oligomers (40)(41). The obtained dependence of the characteristic value of probe density 1/*γ* as a function of *I*_*s*_ is shown in the inset of Figure 6. The values are compatible with an inverse proportionality between *γ* and *I*_*s*_, in agreement with the expected dependence of the free-energy barrier with ionic strength. A deeper insight on the origin of the electrostatic repulsive barrier at the surface functionalized with ssDNA probe is given by the analysis of the dependence of the parameter *γ* on the physical features of the probe layer. According to the HBZ model, the electrostatic penalty *γ* takes the following form (37):

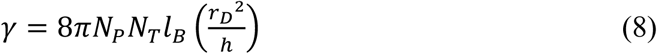

where *N*_*P*_ and *N*_*T*_ are number of bases for probe and target strand respectively, *l*_*B*_ is the Bjerrum length, *r*_*D*_ is the Debye length and *h* is the estimated layer thickness. Given the proportionality of *r*_*D*_ with 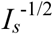 (42), the value of *γ* is expected to scale with 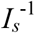, in agreement with the measured values reported in Figure 6 (inset).

## 4 Discussion

### 4.1 Strength of the electrostatic repulsion

The analysis of hybridization kinetics measured by RPI confirms the relevant role of electrostatic repulsion in the observed reduction of hybridization strength on a surface. This effect is ascribed to a free-energy barrier between the free solution state and the bound state of ssDNA targets. In agreement with the VP and HBZ models, the proposed NLER kinetic model adopts a single parameter, *Γ*, to account for such surface repulsion effect. According to our analysis based on Eq. 6, the value of *Γ* can be experimentally extracted through suitably designed experiments, in a range of probe and target lengths, probe surface density, and ionic strength in which the surface repulsion provides a modification of the hybridization kinetics relative to a simple Langmuir model. However, according to the proposed NLER model, the electrostatic free-energy barrier can be relevant even in conditions in which the surface hybridization is indistinguishable from an ideal Langmuir process, hence contributing to the observed weakening of the hybridization strength on a surface (19)(7).

The quantification of the electrostatic repulsive barrier originating at the surface of a DNA biosensor has been addressed in previous works. In (43) it was shown that the data from (19), taken for a 25-mer hybridization at 1 M NaCl, are consistent with a value of *Γ* = 3 at *s*_*p*_ = 10^11^ mm^-2^, whereas a value of about *Γ* = 11.6 would be expected from Eq. 7. In our study, we obtained a value of *Γ* ≈ 1.2 for 12-mer hybridization at 150 mM NaCl and *s*_*p*_ = 10^11^ mm^-2^ (Table 2). Considering only the expected scaling of *Γ* with *N*_*P*_*N*_*T*_ and with 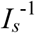 (Eq. 8), which in this case roughly compensate, our values of *Γ* remains from 3 to 10 times smaller than those estimated in (43).

Different hybridization regimes were proposed in (13) depending on the value of strength of the surface repulsion: pseudo-Langmuir (PL), suppressed hybridization (SH), and no hybridization (NH). In (17) it was estimated that for a 20mer directly immobilized on a surface the PL-SH and the SH-NH borders can be placed at *Γ* = 2.5 and *Γ* = 13, respectively. The results of our study are coherent with the conditions between an apparent Langmuir behaviour at small *s*_*p*_ (PL) and a more complex non-Langmuir kinetics (SH), in which the repulsive barrier changes significantly during the hybridization. Therefore, a value of *Γ* around 2.5 would be expected. We explored a range of *s*_*p*_ from 2 to 15·10^10^ mm^-2^, corresponding to a range of *Γ* of about 0.2-1.8 for the no linker and ss-linker probe types (Table 2), thus similar to the estimated threshold, although slightly smaller. A major difference between our experiments and those of (19) and (13) is that we immobilized the DNA probes on a 3D copolymer coating forming a thin hydrogel layer (44), instead of a compact monolayer obtained by direct binding of DNA to the sensing surface. Therefore, from Eq. 8, the apparent discrepancy in the value of *Γ* can be attributed to a larger thickness *h* of the probe surface layer in our case.

### 4.2 Thickness of the surface NA layer

Since all other parameters in Eq. 8 are known or can be easily estimated, we can derive the value of the effective thickness *h* of the region in which the repulsive potential is confined (Figure 5a). In the NLER model, the profile of the repulsive potential along the z coordinate perpendicular to the surface is simply approximated by a step function that remains constant within a thickness *h* and then decreases sharply to the bulk value of the solution. It must be noted that the actual potential will instead change gradually with the distance from the surface (17), hence the parameter *h* represents the effective thickness of the step-like potential providing the same behaviour of the real system. For ionic strengths around physiological conditions, the potential is expected to decrease to the bulk solution value within a few nanometres above the NA layer thickness (24)(51). In contrast, if the NAs are immobilized on a 3D polymer coating, the z-profile of the repulsive potential is expected to be smoother. In the experiments performed in this study, the ssDNA probes were immobilized on the biosensor surface through a multi-functional polymeric coating capable of swelling in aqueous buffer, forming an hydrogel layer with a thickness of about 10 nm when hosting ss or dsDNA (44). Therefore, *h* is expected primarily to depend on the polymer thickness *h*_*c*_ and on the NA layer thickness *h*_*p*_ as *h* = *h*_*c*_ + *h*_*p*_. The characteristic size of the 12mer ssDNA can be estimated assuming a persistence length of about 2.5 nm and a self-avoiding polymer scaling yielding *h*_*p*_ ≈ 5 nm (45), hence *h* is expected to be within 15 nm. In contrast, the value of *h* obtained from Eq. 8 for *N*_*p*_ = *N*_*p*_ =12 is about 125 nm, hence much larger than the expected thickness of the 3D probe layer on the surface. Notably, a similar discrepancy between the measured values of *Γ* and those estimated by Eq. 8 was mentioned also in (43), as discussed above. Here we propose two corrective factors to reconcile the experiments and the theoretical model. A first correction is performed considering that not all the phosphate groups of the ssDNA bring a unitary negative charge. This effect is accounted by the so-called Manning condensation (17) and yields to an effective ssDNA charge of 55% of the fully ionized molecule. Remarkably, in (17) it was reported that this charge renormalization provided the best agreement of a modified Poisson-Boltzmann model with experimental data, hence implying a complete exclusion of mobile counter ions from the DNA surface layer. Since the number of charges enters Eq. 8 through the length of both probe and target DNA, this correction yields a 30% reduction of the calculated *Γ* for a given *h*. A second correction that further reduces the apparent value of *Γ* is obtained considering that in our experiments not all the surface DNA probes are available for hybridization, as indicated by the yield *Ψ* extrapolated from the saturation of the probes at large concentration of target strands. Therefore, only that fraction of probes undergoes a twofold increase of charge, whereas all the probes, not just the hybridized fraction, are responsible for the overall electrostatic repulsion at the surface. From the inspection of Eq. 6 and Eq. 8, a constant additive term in *s*_*p*_ that is not multiplied by (1+*nϕ*) accounts for an increase of the experimentally observed value of *Γ* by a factor 1/*Ψ*, corresponding to a threefold increase for *Ψ* = 30% as for the data in Figure 2. Coherently, the value of *h* in Eq. 8 yielding such larger value of *Γ* is three times smaller. Together with the first correction, an overall reduction of *h* of about a factor of 10 is obtained, hence leading to a thickness of the copolymer layer of *h*_*c*_ = *h* – *h*_*p*_ ≈ 8 nm, in agreement with previous measurements (35)(54). Interestingly, this result suggests that smaller values of *Γ*, hence a reduction of the surface repulsion, can be theoretically obtained for much larger thickness of the 3D functional layer. However, in optical label-free biosensors, distributing the probe molecules at constant *s*_*p*_ along a large thickness can yield to a decrease of signal response upon hybridization, hence an optimal intermediate condition can be preferred.

### 4.3 Origin of the surface weakening of hybridization

The analysis of the hybridization at different surface probe densities enables to extrapolate the expected kinetics and equilibrium strength at very low values of *s*_*p*_, when the repulsive electrostatic barrier vanishes, according to Eq. 7. In this case, the kinetic rate constant for association is given by 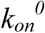, whereas *k*_*off*_ is found to have a much weaker dependence on *s*_*p*_, for the no linker and ss-linker probe types. Accordingly, the dissociation equilibrium constant at very low *s*_*p*_ is given by 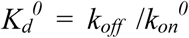. Table 2 reports these values for the studied hybridization schemes. It is interesting to compare the obtained values of 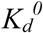 with those for both probe and target ssDNA freely diffusing in solution that can be computed by standard thermodynamic approaches (46). The estimated dissociation constant for the 12mer hybridization in solution remains much lower than 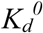 reported in Table 2. Remarkably, as discussed above, a hybridization yield *Ψ* < 1 suggests the presence of a constant additive term in *s*_*p*_, which provides an equivalent correction factor *e*^*Γ*(1-*Ψ*)/*Ψ*^ to the apparent association constant 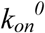 obtained from the fit of the binding curve with Eq. 6. For *Γ* = 1.2 and *Ψ* = 30% this correction factor is more than one order of magnitude, hence leading to values of 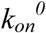 and 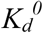 similar to those estimated for hybridization in solution.

Despite the major role of electrostatic repulsion in surface hybridization, other factors can contribute to the weakening of the hybridization strength relative to the same interaction in solution. The possible sources include strand-surface interaction and inter-strand interaction (9). We observed a significant non-Langmuir behaviour even in the case of immobilized strands at a distance larger than their expected lateral occupancy, hence confirming that the origin of the non-Langmuir behaviour is not the inter-strand interaction and that the extrapolation of the hybridization strength at low *s*_*p*_ is not affected by possible inter-strand interaction.

In a previous work we showed that a very weak interaction with the surface can induce a strong decrease of affinity for hybridization because of a simple competition effect (11). This is a peculiar feature of the molecular interaction between complementary NAs, in which the binding sites are spread along the entire molecular length. The temporary unavailability of a single base of the probe strand does not prevent the hybridization but provides a strong effect on hybridization kinetics. Accordingly, on the one side the presence of a polymeric coating with a 3D distribution of conjugation sites can increase the thickness *h* and hence reduce *Γ*, on the other side it can provide more chance of weak interactions, even simply steric, with the immobilized DNA probes, hence reducing the hybridization strength with the target in solution. On the basis of these arguments, an optimal surface functionalization with DNA probes can be achieved combining the ds-linker probe type with a conjugation layer providing suitable thickness, 3D distribution of conjugation sites and minimal interaction with the ssDNA probe.

## 5 Conclusions

The results of this study confirm that the electrostatic repulsion is a major source of the well-known weakening of DNA hybridization on a surface in a wide range of conditions. Despite the strong effect on the equilibrium and kinetics of hybridization, a standard analysis of the binding curves can show only small deviation from an ideal Langmuir behaviour. However, a two-dimensional analysis of the hybridization curves as a function of both *c*_*t*_ and the surface probe density *s*_*p*_ more easily reveals a non-Langmuir dependence coherent with a repulsive potential proportional to the overall density of NA bases on the surface, according to Eq. 6.

The results of this study have direct consequences on the design of DNA arrays. In practice, in the explored conditions the label-free signal due to hybridization is always found to increase with the surface density of probes. Therefore, for the purpose of assay design, larger values of *s*_*p*_ enable to achieve larger signals at equilibrium for any concentration of target *c*_*t*_. However, the kinetics of hybridization can be strongly reduced at large *s*_*p*_ by two phenomena: the surface electrostatic repulsion and the crowding of immobilized probes. The latter effect only occurs for large enough fractional coverage *ϕ* of probes, hence typically close to saturation, whereas the electrostatic penalty can be effective at any values of *ϕ* and *c*_*t*_ and directly contributes to reduce the observed equilibrium constant for surface hybridization. Accordingly, a correct absolute quantification of target concentration derived from the assay response should necessarily account for the weakening and slowing down of hybridization, which both depend on the surface density of probes.

Interestingly, the large net charge of NA can be considered as functional to preserve a large specificity of hybridization even at large concentrations. The electrostatic repulsion between two NA strands in solution effectively increases the threshold of the attractive strength required to form a stably paired complex, hence the minimum number of consecutive complementary bases. Indeed, uncharged DNA mimics such as peptide nucleic acids (PNA) or phosphorodiamidate Morpholino oligomer (PMO), although they may provide larger affinities for hybridization with DNA in controlled conditions, they typically also display lower solubility and larger non-specific binding that brings to relevant background signals when used in assays applications (8). Analogously, it can be argued that the enhanced repulsion originating in surface-based DNA biosensors favours the specificity of molecular recognition at the cost of sensitivity, relative to DNA probes freely diffusing in solution. From the results of this study, we can estimate a DNA concentration in solution, at which the electrostatic repulsion starts inducing non-negligible effects on the hybridization kinetics. From Eq. 6, we can assume that the hybridization behaviour deviates from a Langmuir model for *Γ* > 0.1. This corresponds to *s*_*p*_ of the order of 10^10^ mm^−2^ for the strand probes used in this work. Considering a 3D distribution of the probes over a thickness of about 13 nm, the corresponding volume density is about 8·10^17^ molecules in 1 ml, or 5 mg/ml for a 12-mer DNA probe. As a comparison, the average concentration of DNA within the nucleus is of the order of 10 mg/ml (47), with large density fluctuations in space. Therefore, the conditions achieved on the surface of DNA biosensors and the corresponding effects on hybridization can be rather common in nature and can play a biological role in the cellular nucleus. Overall, the kinetic modelling of these elementary DNA-based interactions is expected to guide the design of more complex functional structures immobilized on a surface and provide a pathway for kinetic optimization of DNA nano-machines.

## Author contributions

L. V. designed research, analyzed data and wrote the manuscript, T. C. and L. C. performed research, M. S. contributed analytic tool, M. C. designed research, G. Z. and M. B. designed research and wrote the manuscript.

## Acknowledgements

This work has received funding from Regione Lombardia and FESR, Linea Accordi per la Ricerca, through the “NeOn” project (ID 239047), and from Ministero dell’Istruzione, dell’Università e della Ricerca, PRIN2017, through the “Soft Adaptive Networks” project. We thank ProXentia Srl (Milano, Italy) for providing the RPI sensing cartridges. MB is a founder and shareholder of ProXentia Srl.

